# An inter-channel cooperative mechanism mediates PIEZO1’s exquisite mechanosensitivity

**DOI:** 10.1101/2021.04.16.440217

**Authors:** Tharaka Wijerathne, Alper D. Ozkan, Wenjuan Jiang, Yun Luo, Jérôme J. Lacroix

**Affiliations:** Graduate College of Biomedical Sciences, Western University of Health Sciences, 302 E 2nd st, Pomona, CA 91709, USA; College of Pharmacy, Western University of Health Sciences, 302 E 2nd st, Pomona, CA 91709, USA

## Abstract

The bowl-shaped structure of PIEZO channels is predicted to flatten in response to mechanical stimuli, gating their pore open. However, how this unique structure allows them to detect exquisitely small changes in membrane tension remains unclear. Here, using pressure clamp electrophysiology, modeling, and molecular dynamics simulations, we show that the single channel open probability of PIEZO1 increases weakly with respect to pressure-induced tension. In contrast, when multiple channels are present in a membrane patch, channel open probability increases steeply as a function of the number of open channels. These cooperative effects are consistent with an inter-channel energetic repulsion due to the local membrane deformation created by the non-planar PIEZO structure. When channels open, this deformation shrinks, allowing open channels to diffuse closer to each other, thus delaying closure. This study reveals how PIEZO1 channels acquire their exceptional mechanosensitivity and suggests a possible mechanism by which cells could rapidly tune mechanosensitivity.

## INTRODUCTION

Mechanosensitive PIEZO1 and PIEZO2 channels contribute to an astonishing diversity of mechanosensory processes across most physiological systems^1^. PIEZO1 channels are directly sensitive to physical deformations of the lipid bilayer and thus do not require cellular components other than the cell membrane to sense mechanical forces^2^. PIEZO1’s mechanosensitivity is best quantified using the pressure-clamp electrophysiology technique in which a membrane patch in a cell-attached mode is stretched by application of positive or negative pressure to the backside of the recording pipette. Using this technique, the membrane tension necessary to open PIEZO1 has been reported to be lower compared to other known mechanosensitive channels^3–9^. What mechanism confer PIEZO1 channels their exquisite mechanosensitivity?

The closed structure of homotrimeric PIEZO channels consists of three spiraling peripheral domains arranged around a central pore, defining a unique bowl-like architecture^10–13^. A prevailing gating mechanism posits that membrane stretch increases channel open probability by promoting a flatter channel conformation^13,14^. Owing to its unique bowl shape, the closed PIEZO structure is predicted to impose a large deflection to the surrounding lipid bilayer called PIEZO footprint^15^. The membrane deformation energy cost associated with the PIEZO footprint brings about work to flatten the bowl-shaped channel: as tension increases, so does this energy, potentially converting lateral membrane tension into flattening gating motion.

The footprint-based gating paradigm predicts PIEZO channels sense both membrane tension and curvature^15^. This idea is supported by two independent experimental results. First, PIEZO1 channels reconstituted into liposomes are more curved in smaller liposomes than they are in larger ones^14^. Second, PIEZO1 channels spontaneously open in absence of external force when reconstituted into asymmetric (non-flat) artificial membranes, but not into symmetric (flat) ones^2,16^.

The sensitivity of PIEZO channels to the surrounding membrane curvature has surprising consequences. The overlap of adjacent PIEZO footprints would necessarily increase membrane deformation energy and thus is accompanied by an energy penalty. Unless sufficient energy is provided to overcome this penalty, neighboring channels are thus predicted to remain at bay^15^. If adjacent channels were allowed to move near each other and overlap their footprints, the increased membrane deformation energy would bring about tension-independent work to flatten them and increase open probability. Such a cooperative gating phenomenon has been observed in Molecular Dynamics (MD) simulations in which periodically mirrored PIEZO1 channels, virtually brought near each other by reducing the spatial dimensions of the simulated system, overlap their footprints and spontaneously flatten, enabling their pore to open^17^.

In this study, we sought to determine whether such cooperative effects occur in living cells. Using pressure-clamp electrophysiology, we show that single PIEZO1 channels are weakly mechanosensitive but dramatically increase open probability as more channels open in multichannel recordings, indicating strong cooperative interactions. Because the size of the PIEZO footprint depends on channel curvature, the footprint of flatter open channels is predicted to be smaller than that of more curved closed channels. Thus, PIEZO1 channels should diffuse closer to each other when they are open, as confirmed here by MD simulations. When open channels diffuse near each other, their closure would instantly extend their footprints, incurring a footprint overlap energy penalty. Open channels are thus anticipated to remain open for longer periods of time as more open channels diffuse around them, consistent with our observation that the closure rate decreases as the maximal amplitude of PIEZO1-mediated currents increases.

## RESULTS

### Single PIEZO1 channels are weakly mechanosensitive

To test whether PIEZO1 gating is influenced by intermolecular cooperative effects, we transfected HEK293T^ΔP1^ cells (where endogenous PIEZO1 expression is abolished^18^) with a plasmid encoding mouse PIEZO1 and measured mechanically-evoked currents in these cells using the pressure clamp technique in the cell-attached configuration. While patches from untransfected cells did not produce mechanically-evoked currents, patches from transfected cells yielded robust pressure-dependent inward currents. By incrementally increasing the amplitude of pipette suction pulse (negative pressurization), the amplitude of these currents saturates (Figure 1a). We noticed that when the saturating currents were large, they tend to decay exponentially, a hallmark of PIEZO1 channel inactivation. However, when these currents were small, inactivation was slower (Supplementary Figure 1). Plotting the relative peak current (before inactivation sets), 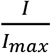,as a function of the pressure pulse, *p*, produces sigmoid-like curves that tend to be shifted toward more negative pressures as *I*_*max*_ decreases (Figure 1b). These sigmoid curves can be well fitted using a classical Boltzmann equation (see Supplementary Appendix 1):

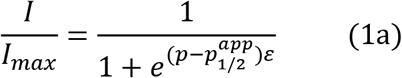

**Figure 1.**
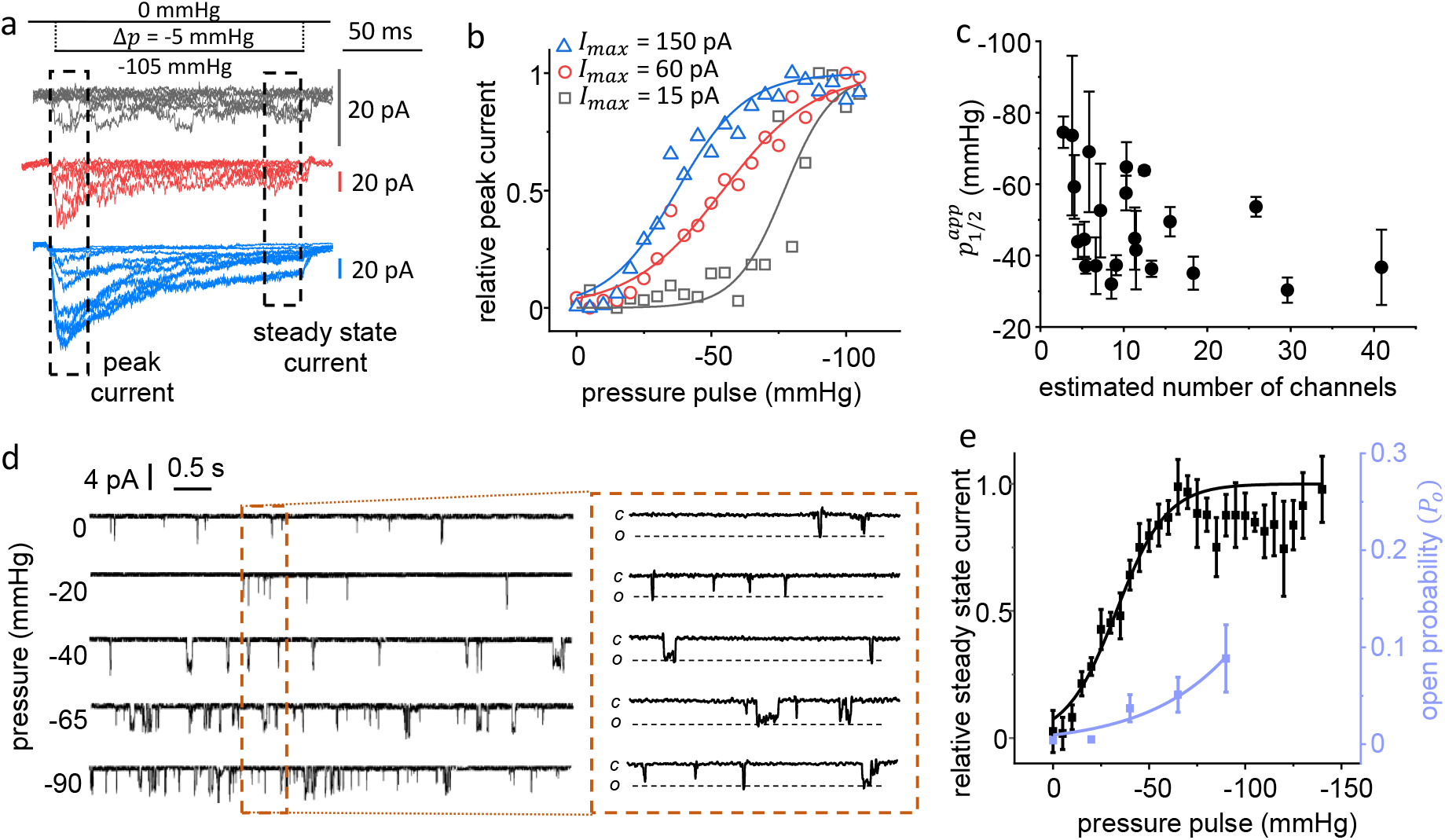
PIEZO1’s mechanosensitivity depends on the number of channels in a patch. (**a**) Examples of pressure-elicited PIEZO1 currents in patch with varying peak current amplitude. V = −100 mV (**b**) Relative peak current from data in (a) plotted as a function of pulse pressure. (**c**) The apparent half-activation pressure obtained from individual patches is plotted as a function of the estimated number of active channels in the patch. Error bars = standard deviation from fit with equation (1a). (**d**) Example of 20 sec-long single channel recordings obtained at the indicated steady state pressure. 500 ms snippets from each trace are shown in the insert. (**e**) Comparison of the PIEZO1 pressure activation curve obtained in steady state conditions for single channel patches (violet circles; number of independent patches/pressure in mmHg: 2/0, 6/−20, 5/−40, 4/−65, and 4/−90) and for macroscopic patches (black circles, 11 independent patches). Error bars = s.e.m.

With 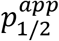 the apparent pressure value producing half of the maximal current 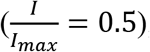, and *ε*, a slope factor. We compiled 22 independent macroscopic recordings, fitted the corresponding 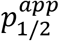 values, and plotted them as a function of the estimated number of channels in the patch.

The number was obtained by divided *I*_*max*_ by the single channel current amplitude. In our experimental conditions (−100 mV transmembrane voltage and 140 mM KCl pipette solution), the unitary current was ≈ 5.5 pA. This plot shows that, as more channels populate the patch, the fitted 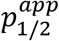 values tend to be less negative (Figure 1c). This suggests that large PIEZO1 channel populations have a lower activation threshold compared to smaller channel populations.

To determine the inherent mechanosensitivity of single PIEZO1 channels, we next obtained > 60 sec-long single channel recordings from patches maintained at a steady-state negative pressure (Figure 1d). The dwell times of opening and closure events follow mono or dual gaussian distributions when plotted in a logarithmic time scale, indicating the duration of our recordings was long enough to capture ensembles of stochastic gating events (Supplementary Figure 2a). To rule out the presence of multiple channels, we applied a strong pressure pulse at the end of each record and discarded those containing more than one conductance level (Supplementary Figure 2b). In agreement with our observation that PIEZO1-mediated currents do not decay when their saturating amplitude is low, single PIEZO1 channels did not exhibit time-dependent loss of activity across the length of our recordings in our experimental conditions. The single channel open probability, *P*_*o*_, was very low and only increased from ≈ 0.005 to ≈ 0.1 when pressure decreased from 0 to −90 mmHg (Figure 1e). Sustained patch pressurization below −90 mmHg induced electrical instability and membrane rupture, precluding accurate *P*_*o*_ determination at more negative pressures. We fitted the *P*_*o*_ vs. *p* plot using the following Boltzmann equation (see Appendix 1):

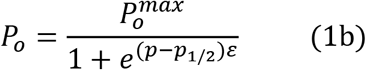

Because *P*_*o*_ does not saturate between −65 and −90 mmHg, our data does not allow us to reliably estimate 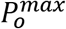. Although increasing tension is predicted to promote an open state by gradually stabilize a flat channel conformation, it is unclear whether the flat conformation is associated with an open probability of 1. Assuming 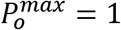, fitting our single channel data with equation (1b) yields *p*_1/2_ = −178 ± 19 mmHg and *ε* = 0.026 ± 0.006 mmHg^−1^ (R^2^ = 0.946). Assuming 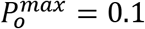 (the minimum possible value according to our data) the fit yields *p*_1/2_ = −58 ± 5 mmHg and *ε* = 0.053 ± 0.011 mmHg^−1^ (R^2^ = 0.957) (Table 1). To compare single channel vs. macroscopic data in similar steady-state conditions, we plotted the relative steady-state current, 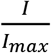, obtained from macroscopic patches with at least ≈ 20 channels (*I*_*max*_ < −100 pA) as a function of pressure and fitted this trace with equation (1a) (Figure 1e). The fit yields 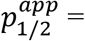 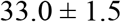 and *ε* = 0.077 ± 0.009 mmHg^−1^ (R^2^ = 0.927). For comparison, when 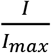 is taken at the peak current, 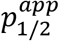 was slightly lower (−25 ± 2 mmHg) and *ε*, slightly larger (0.117 ± 0.018 mmHg^−1^) (Supplementary Figure 3). Under the 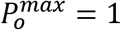 assumption, our data show that the pressure necessary to elicit half of the maximal current for a large channel population decreases by about 6-fold and the slope factor of the pressure-activation curve increases by about 3-fold compared to single channels. Under the 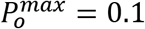 assumption, our data still suggest the PIEZO1 mechanosensitivity is weaker for single channel (*p*_1/2_ shifted towards more negative values and smaller *ε*) compared to channel populations, albeit to a lower extent.

**Table 1:** Single channel parameters obtained by fitting the single channel pressure-activation curve (Figure 1e) using equation (1b) and assuming different 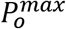 values.

| $P_o^{max}$ | $P_{1/2} \text{ (mmHg)}$ | $\varepsilon \text{ (mmHg}^{-1}\text{)}$ | $R^2$ |
| --- | --- | --- | --- |
| 1 | $-178 \pm 19$ | $0.026 \pm 0.006$ | 0.946 |
| 0.9 | $-174 \pm 19$ | $0.026 \pm 0.005$ | 0.946 |
| 0.8 | $-168 \pm 17$ | $0.026 \pm 0.005$ | 0.946 |
| 0.7 | $-161 \pm 16$ | $0.027 \pm 0.005$ | 0.947 |
| 0.6 | $-154 \pm 15$ | $0.027 \pm 0.005$ | 0.947 |
| 0.5 | $-145 \pm 13$ | $0.028 \pm 0.005$ | 0.948 |
| 0.4 | $-133.5 \pm 11$ | $0.028 \pm 0.005$ | 0.949 |
| 0.3 | $-118 \pm 9$ | $0.030 \pm 0.005$ | 0.951 |
| 0.2 | $-96 \pm 6$ | $0.033 \pm 0.006$ | 0.956 |
| 0.1 | $-58 \pm 5$ | $0.053 \pm 0.011$ | 0.957 |

### PIEZO1 channel populations do not gate independently

To further explore how modulating the number of channels affects their mechanosensitivity, we obtained steady-state current traces exhibiting more than one open conductance levels (Figure 2a). We estimated the total number of channels, *n*, by taking the maximal conductance level observed during at least 60 seconds of continuous recording. Since the inherently large electrical noise of large channel populations reduce the accuracy of our estimates of *n*, we arbitrarily selected traces with *n* < 15. Although most patches harbored many more channels, we were able to collect a total of 63 multichannel records with 2 < *n* < 15 at 3 distinct patch pressures (−20, − 40, and −90 mmHg). Since it was not possible to predict the number of channels per patch, we were not able to obtain all possible *n* values at all tested pressures or to obtain replicates for some combinations of patch pressure and channel number. In our multichannel traces, the dwell time of each conductance level plotted on a logarithmic time-axis display a bell-shape resembling a gaussian distribution, suggesting our sampling duration was long enough for meaningful statistical analysis (Supplementary Figure 4a). Regardless of patch pressure, we found that the maximal number of channels did not increase when applying a strong test pulse at the end of our recordings, further validating our estimation of the total number of channels (Supplementary Figure 4b). In agreement with previous observations, channel activity did not diminish over time, supporting the notion that inactivation does not appear to take place when few channels are present in the patch (Supplementary Figure 5).

**Figure 2.**
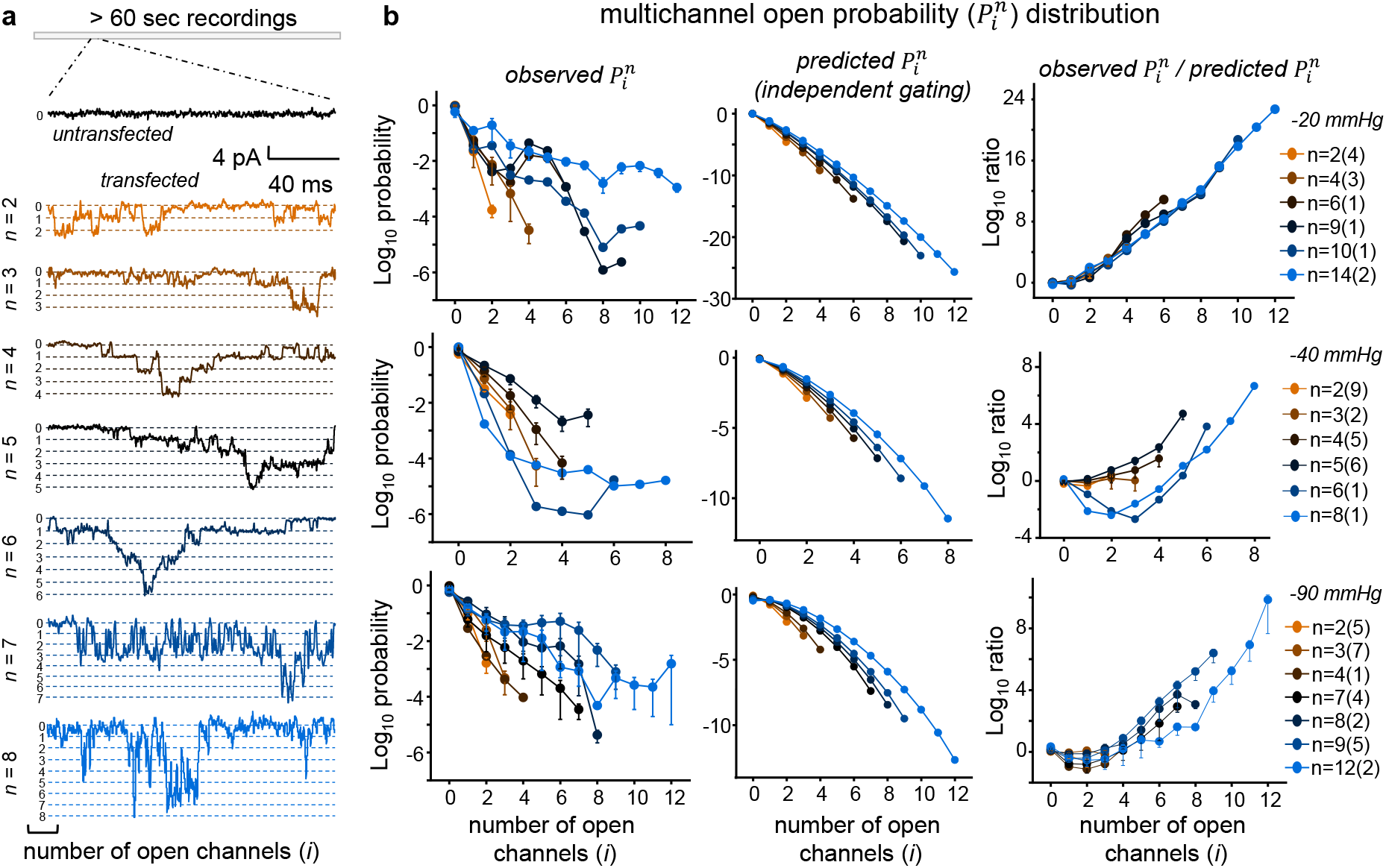
PIEZO1 channels do not gate independently in multichannel patches. (**a**) Representative snippets from multichannel current traces from patches pressurized at −40 mmHg and containing a variable number of channels (V = −100 mv). (**b**) The observed probability of *i* channels to be open in a patch containing *n* channels 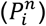 is compared with the 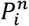 predicted by equation (2) for independent channels. In panels (a) and (b), the color gradient represents the number of channels in the patch. In the legend of panel (b), the number in parentheses indicate the number of independent patches for each n value. Error bars = s.e.m. Continuous lines connecting discrete probability values have no physical meaning and are displayed for clarity only.

For each tested pressure, an event extraction analysis allowed us to determine 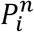, the probability of *i* channels being open in a patch containing *n* channels, as a function of *i* (see Methods). If PIEZO1 channels were gating independently, the observed 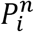 would follow the combinatorial equation (see Supplementary Appendix 2.1):

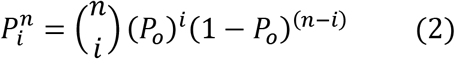

Using *P*_*o*_ values from Figure 1e, equation (2) poorly predicts the observed 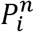 (Figure 2b). The discrepancy between the 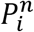 observed from multichannel recordings and those calculated under the assumption of independent gating can be visualized by plotting the ratio of observed vs. predicted 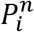: for all tested pressures, this ratio tends to increase in a logarithmic manner by many orders of magnitude as the value of *i* increases, regardless of the total number of channels present in the patch (Figure 2b). By contrast, the 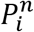 obtained from MATLAB simulations in which multiple channels gate independently are well described by equation (2) (Supplementary Figure 6).

### Modelization of inter-channel cooperativity

For simplification, we postulate that the discrete gating transitions observed in multichannel patches are predominantly mediated by transitions between two states, closed (C) and open (O):

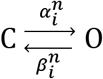

with 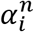 and 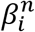, respectively the microscopic opening and closure rate in patches containing *i* open channels and *n* total channels. Since the 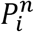 values appear to increase as a function of the number of open channels in the patch, we hypothesize that the thermodynamic tchoensctlaonsted/roivpienngequilibrium, 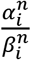 scales by a constant parameter, *k*, for each iteration of *i*:

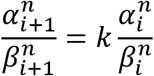

This sequential cooperative model leads to the following expression to predict 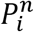 (see Supplementary Appendix 2.1):

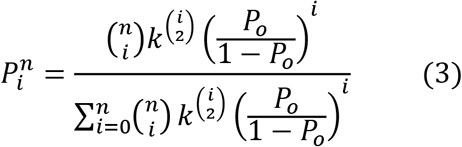

In this model, inter-channel cooperativity is positive when *k* > 1 and negative when 0 < *k* < 1. When *k* = 1, equation (3) is mathematically equivalent to equation (2) for independent channels (see Supplementary Appendix 2.1). It is noteworthy that the *k* exponent in equation (3) is a binomial coefficient that represents the number of combinations of pairs of open channels. Since *P*_*o*_ and *n* are experimentally known from our single and multichannel recordings, the only unknown fitted parameter in equation (3) is *k*. We log-transformed the 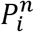 data and calculated the Mean Absolute Error (MAE) between observed 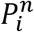 and the 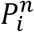 values calculated using equation (3) by varying *k* (see Methods). MAE minimization yielded optimal *k* values of about 2.25, 1.65, and 1.31 for patch pressures of −20, −40, and −90 mmHg, respectively (Figure 3b and Supplementary Figure 7). Plotting the ratio between observed and fitted 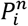 as a function of *I* shows our sequential model fits well our data at −20 and −90 mmHg, as evidenced by the convergence of these ratios towards unity along the *i* dimension (Figure 3b). This convergence was not as good for patches at −40 mmHg, presumably due to noisier data.

**Figure 3.**
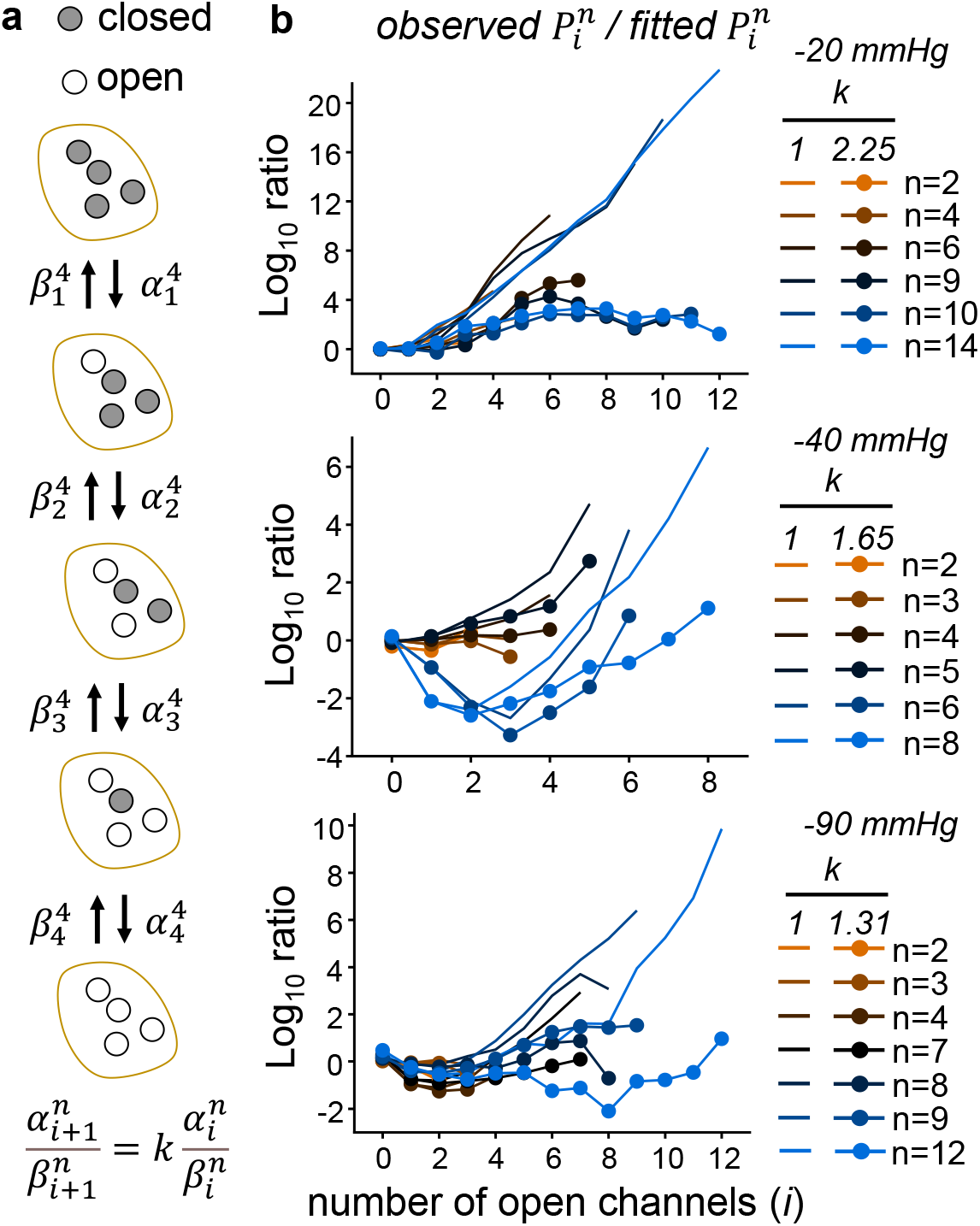
Predicting PIEZO1 multichannel open probabilities with a sequential cooperativity gating model. (**a**) Schematic illustration of the gating model in which *k* represents the cooperativity parameter. (**b**) The ratio of observed vs. fitted mean 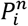 values is plotted for each tested pressure (lines with full circles). These plots are compared with those obtained assuming no cooperativity (*k* = 1, continuous lines). Data and coloring methods are from Figure 2.

We additionally tested two variants of this model. The first one states that the PIEZO1 activation constant, 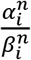, scales by *k* for each iteration of *n* instead of *i*. In this “numeral” model, the exponent of the *k* parameter equals *i*(*n*−1) (Supplementary Appendix 2.2). In the second variant, the exponent of the *k* parameter was arbitrarily set to *i*(2*n*−*i*−1)/2, i.e. the number of combinations of channel pairs in which at least one channel is open. MAE minimization shows that the sequential model produces slightly smaller errors compared to these variants (Table 2). In addition, the shape of the observed 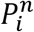 distribution (Figure 2b) more closely resembles the U-shaped distribution predicted by the sequential model rather than the rainbow-shaped distribution predicted by the two other models (Supplementary Figure 8).

**Table 2:** Comparison of fitted *k* values and minimal MAE for the three cooperativity models tested in this study.

| exponent of $k$ parameter | patch pressure (mmHg) | fitted $k$ value | MAE |
| --- | --- | --- | --- |
| $\binom{i}{2}$; equation (3) | -20 | $2.25 \pm 0.055$ | 1.64 |
| | -40 | $1.65 \pm 0.080$ | 0.84 |
| | -90 | $1.31 \pm 0.018$ | 0.89 |
| $i(n-1)$ | -20 | $1.48 \pm 0.012$ | 1.72 |
| | -40 | $1.24 \pm 0.027$ | 0.92 |
| | -90 | $1.05 \pm 0.007$ | 1.05 |
| $\frac{i(2n-i-1)}{2}$ | -20 | $2.00 \pm 0.031$ | 1.79 |
| | -40 | $1.20 \pm 0.047$ | 0.98 |
| | -90 | $1.15 \pm 0.014$ | 1.21 |

### Estimating the number of cooperating channels

How many channels energetically cooperate, on average, in macroscopic recordings? Our sequential model relates this number, *n*, to the relative macroscopic current elicited upon acute pressure stimulation (see Appendix 3):

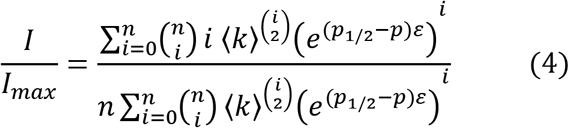

The values of *p*_1/2_ and *ε* depend on 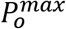 and are obtained from fitting the single channel pressure-activation curve using equation (1b) (Figure 1e). Because *k* varies with *p*, we averaged *k* values fitted from multichannel records at all tested pressures (〈*k*〉). Accordingly, the only unknown parameter in equation (4) is *n*. We compiled a range of macroscopic pressure-activation curves for PIEZO1 based on our data and that from others. These curves are described by equation (1a) with 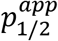 ranging from −25 to −40 mmHg and *ε* ranging from 0.08 to 0.1^2–4,19–21^. These experimental pressure-activation curves are consistent with those predicted by equation (4) with *n* values equal to 14-15 assuming 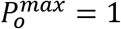, 11-13 assuming 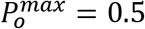, 9 assuming 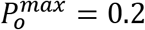, and 7 assuming 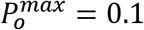 (Figure 4). Interestingly, the pressure-activation curve predicted by equation (4) with 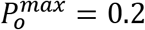 and *n* = 9 seems to best overlap with the consensus experimental curve.

**Figure 4.**
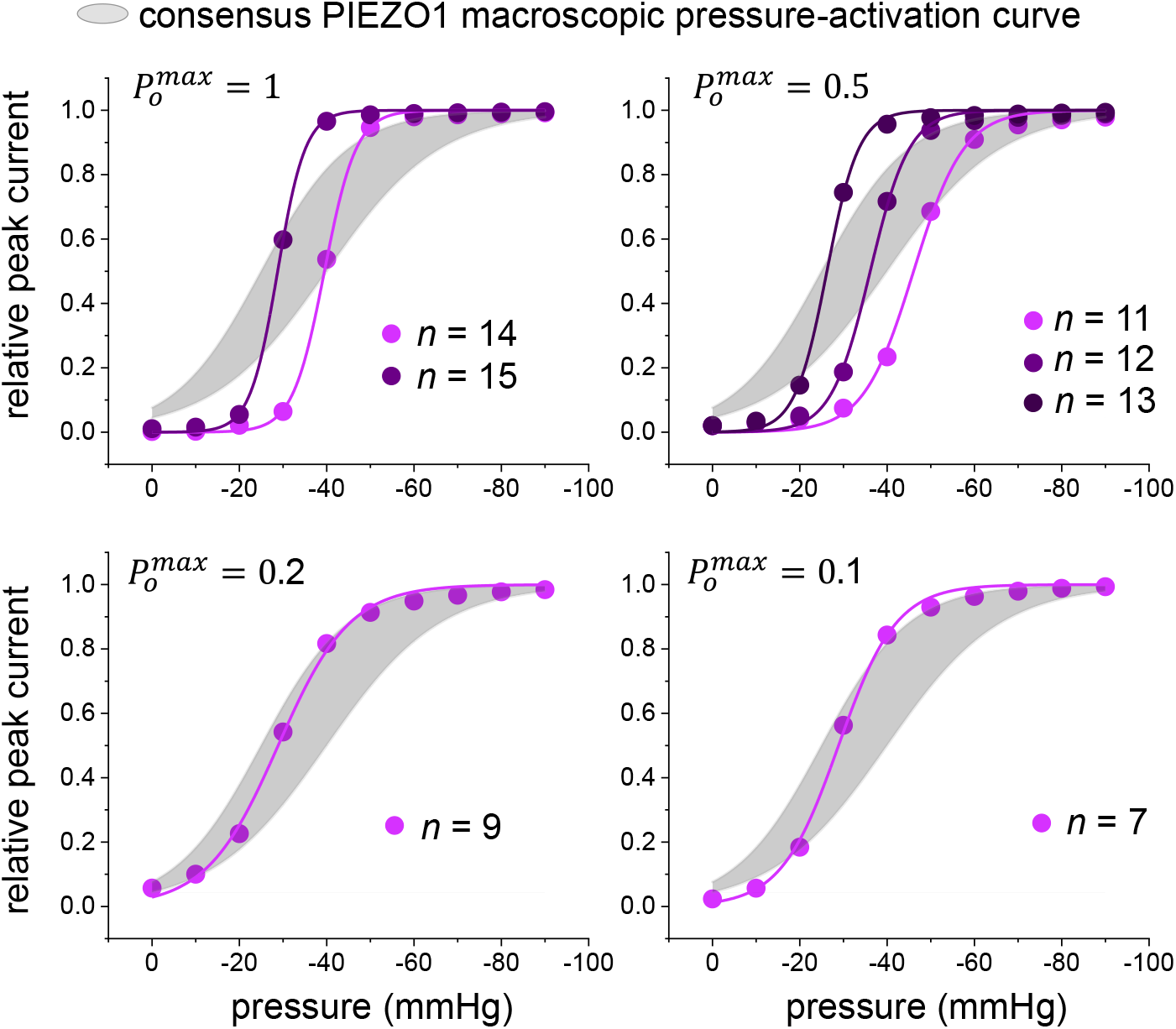
Estimation of the number of cooperating PIEZO1 channels in macroscopic patches. The grey areas represents a consensus pressure-activation curve obtained from independent studies (see text). Colored lines represent hypothetical pressure-activation curves obtained using equation (4) that more closely overlap with the consensus curve assuming different values of 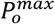. In each case, the fitted discrete number of cooperating channels (*n*) is indicated.

### A diffusion-based mechanism for PIEZO1 cooperativity

The seemingly complex inter-channel cooperativity behavior of PIEZO1 channels can be explained by a simple mechanism based on channel diffusion and membrane energetics. As mentioned earlier, the PIEZO1 gating free energy increases if adjacent channels overlap their footprints. However, the First Law of Thermodynamics posits that this added energy constitutes a penalty. Therefore, the footprint overlap is energetically unfavorable unless this penalty is paid for by another energy source. This energy source could originate, for example, from inter-channel collisions, from the motions of molecular motors tethered to the channels *via* cytoskeletal elements^22^, or from the segregation of channels into lipid microdomains^23^. In such cases, however, the cooperative effects would increase as a function of the total number of channels, not as a function of the number of open channels. What mechanism could thus mediate the cooperative behavior of PIEZO1 channels without utilizing the additional gating free energy brought about by overlapping adjacent footprints?

When PIEZO channels open, their structure is predicted to adopt a flatter conformation. The footprint of open channels is therefore predicted to be smaller than that of closed channels, enabling open channels to diffuse closer to each other. When open channels are near each other, channel closure would return the channel to a curved shape, instantly extending the channel footprint. This footprint extension would likely overlap with that of adjacent channels, instantly incurring the footprint overlap energy penalty. When open channels diffuse near each other, their closure is thus expected to become thermodynamically unfavorable, enabling clustered open channels to remain open for longer periods of time compared to their isolated counterparts (Figure 5).

**Figure 5.**
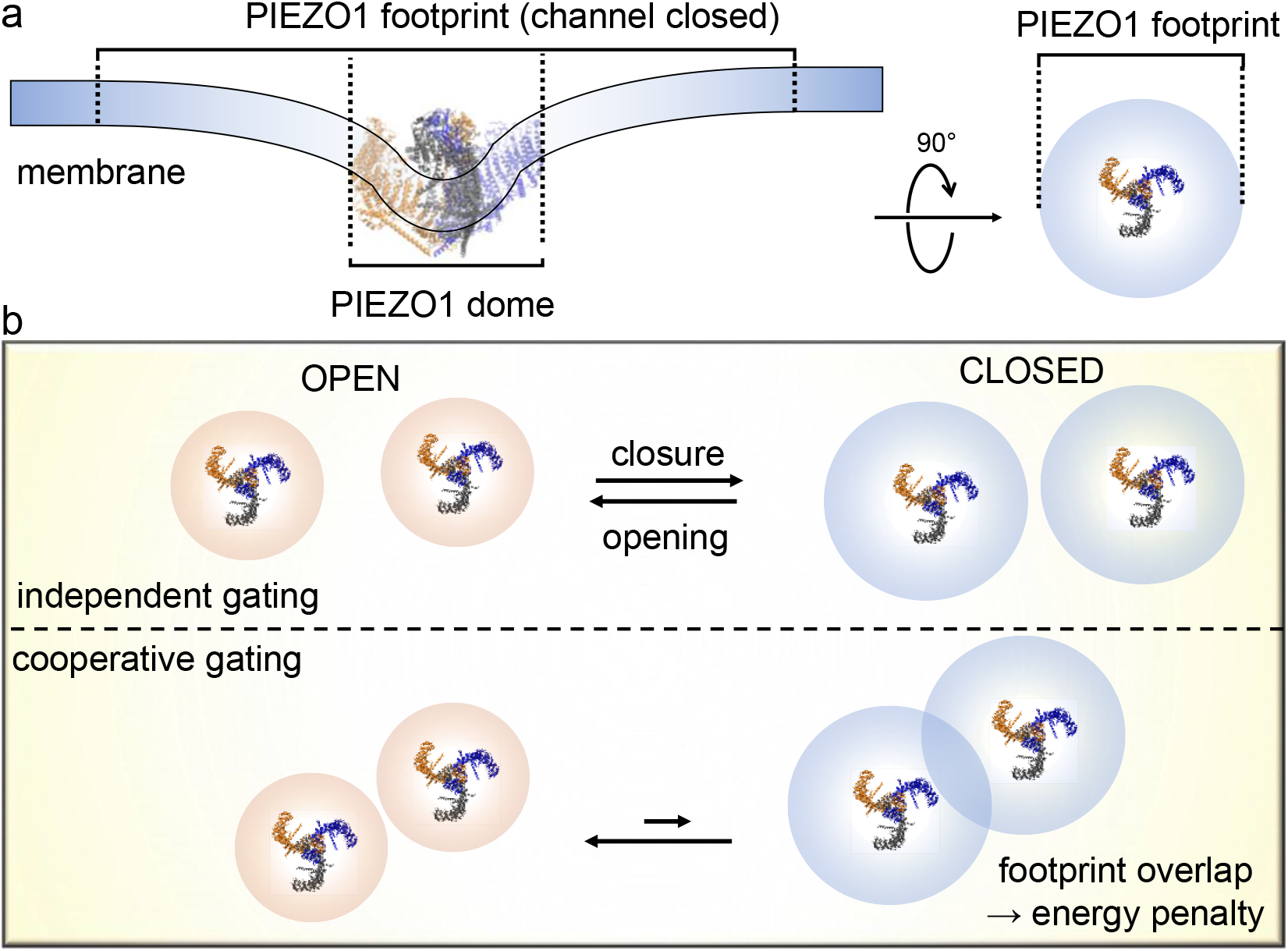
Proposed inter-channel cooperative gating mechanism for PIEZO1. (**a**) Schematic illustration of the PIEZO1 footprint and dome for a closed channel. (**b**) Due to their flatter conformation, open PIEZO1 channels produce smaller footprints. When open channels are near each other, their closure would overlap their footprints, incurring an energy penalty.

### Validation of the proposed cooperativity mechanism

A central tenet of our cooperative mechanism states that PIEZO1 channels diffuse closer to each other when they are open as compared to when they are closed. To test this assumption, we used coarse-grained (CG) simulations to determine the minimal inter-channel distance between pairs of open or closed channels diffusing for ≈ 7 μs in a 52 x 104 nm membrane at 313 or 340 K. The spatial coordinates corresponding to the open and closed conformations were obtained from a recent computational study^17^. In CG simulations, the protein backbone is rigid, but proteins freely rotate and diffuse relative to lipid and solvent molecules. In our case, due to periodic boundary conditions, the minimal inter-channel distance fluctuates between the two simulated channels, either within the same box or between mirrored boxes (Figure 6a). At both temperatures, the minimal pore-pore distance between closed channels remain > 40 nm during the entire trajectory, while for open channels, this distance decreased by 20~30 nm over the same duration (Figure 6c and Supplementary Figure 9). These results are consistent with a long-range footprint overlap energy penalty preventing closed channels to get closer through random diffusion.

**Figure 6.**
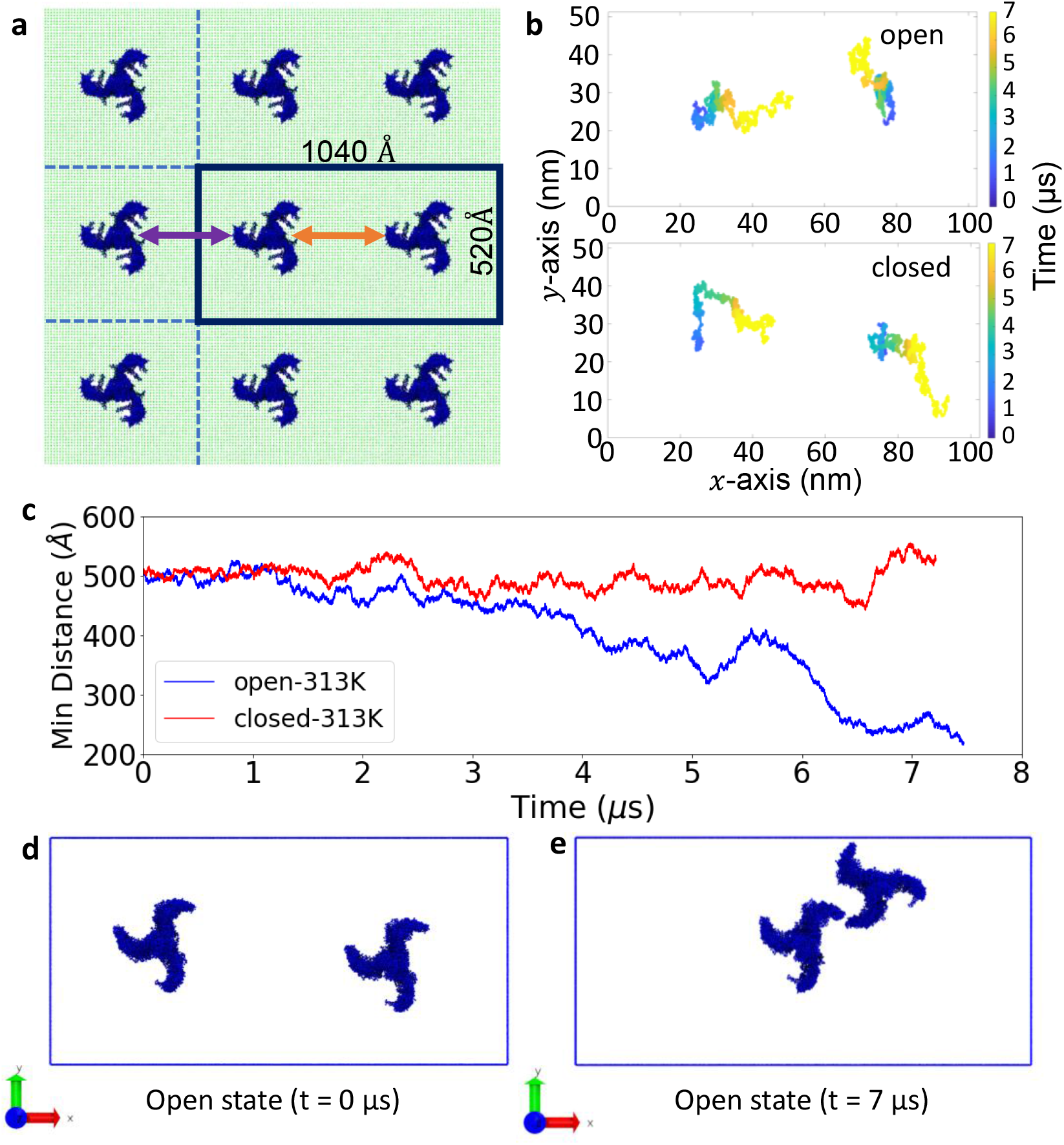
PIEZO1 channels diffuse closer when they are open in coarse-grained MD simulations. **(a)** A pair of closed channels is shown in the simulated box (delineated by the black perimeter) and in surrounding periodic images. Arrows indicate minimum pore-pore distance between one channel and its counterpart in the same (orange) or a mirrored (purple) box. (**b**) Temporally-colored trajectories of the diffusion of the center of mass of each closed channel. (**c**) Time course of minimum pore-pore distance between pairs of open (blue) or closed (red) channels at 313 K. Snapshots of the open channels simulation are shown at the beginning (**d**) or at the end (**e**) of the simulation.

Besides inter-channel distance, our mechanism predicts that increasing the number of open channels would reduce the rate of channel closure, without modulating the rate of channel opening. To test these predictions, we evoked PIEZO-mediated macroscopic currents using a saturating −100 mmHg pressure pulse of a short duration to prevent the development of inactivation. We determined the time course of activation/deactivation by fitting the rising/decaying phase of current traces upon application/removal of the pressure stimulus with a mono-exponential function. Plotting activation and deactivation time constants as a function of the absolute amplitude of the peak current, |*I*_*max*_|, seems to confirm our mechanism. As |*I*_*max*_| increases, channel closure drastically slows down while channel opening accelerates only slightly. Pearson’s correlation coefficient between |*I*_*max*_| and closure time constant is ≈ 0.76, while Pearson’s correlation coefficient between opening time constant and |*I*_*max*_| is ≈ −0.44. A linear regression analysis further shows that, for the same increase of |*I_max_*|, the slowing down of channel closure is about 10-fold larger than the acceleration of channel opening (linear slope of 6.42 ± 1.12 ms nA^−1^ vs. −0.61 ± 2.79 ms nA^−1^) (Figure 7).

**Figure 7.**
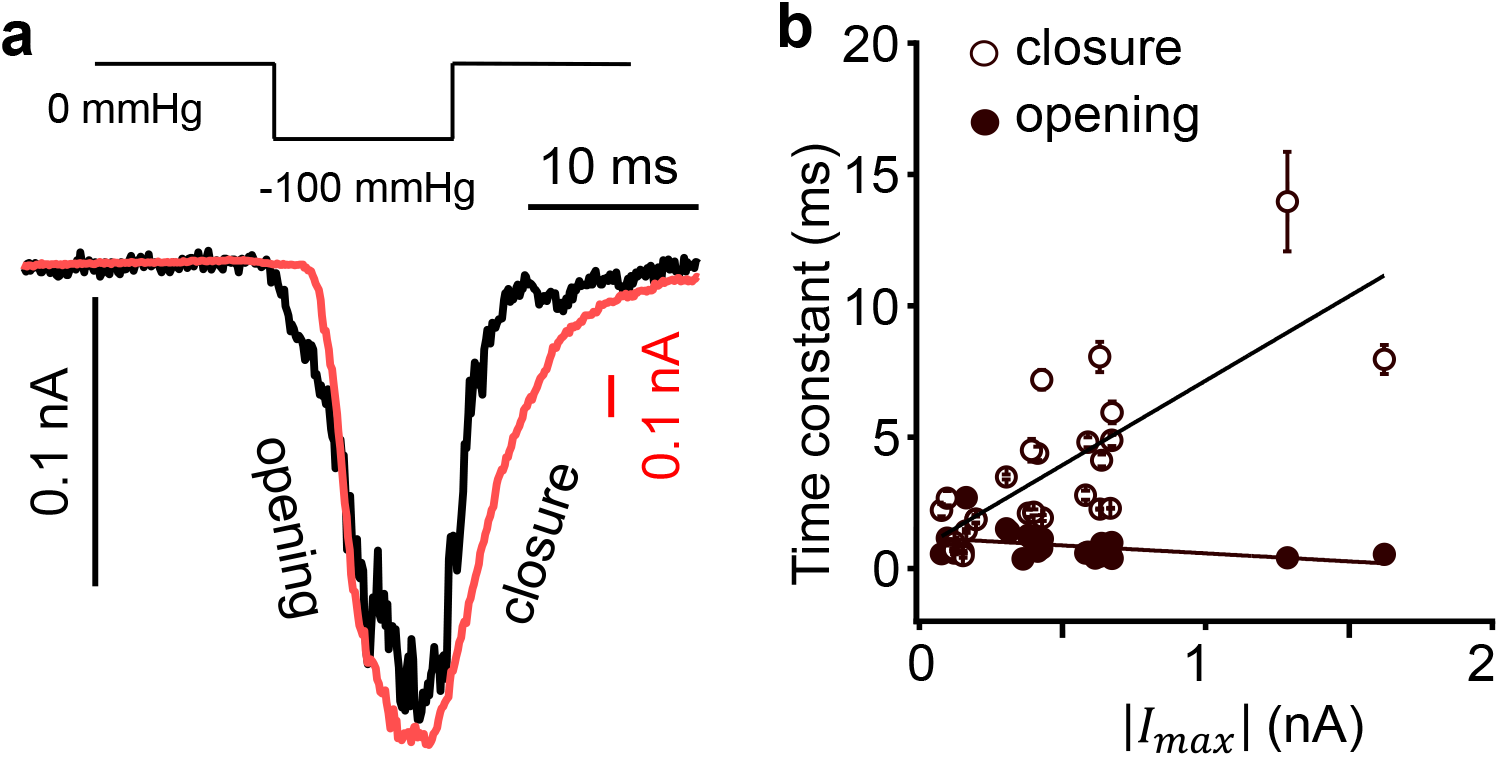
PIEZO1 channels close more slowly when more channels open in the patch. (**a**) Example of macroscopic PIEZO1 current traces of different amplitude obtained with a 10 ms saturating pressure pulse. V = −100 mV. (**b**) The time constant of activation (opening, open circles) and deactivation (closure, filled circles) is plotted as a function of the absolute value of maximal current amplitude (|*I_max_*|). Circles represent independent patches. Error bars = s.e.m. from exponential fit. Lines represent linear fits to the data using y = a + bx. Fitted parameters: a = 1.19 ± 0.16 ms (activation) and 0.74 ± 0.66 ms (deactivation); and b = −0.61 ± 2.79 ms nA^−1^ (activation) and 6.42 ± 1.12 ms nA^−1^ (deactivation).

Finally, our proposed mechanism postulates that the strength of cooperative effects is proportional to the energy penalty that would be incurred if adjacent channels were to overlap their footprints. In a channel cluster, this energy penalty should depend on both the number and surface area of individual PIEZO footprints. Remarkably, the decay length of individual footprint is predicted to shrink as membrane tension increases^15^. Thus, for the same number of open channels, increasing membrane patch pressure is predicted to reduce the total gating free energy contributed by inter-channel cooperativity. Consequently, the cooperative effects are anticipated to decrease as tension increases. This is well consistent with our observation that the value of the fitted *k* parameter decreases with increasing patch pressure, from ≈ 2.25 at −20 mmHg to ≈ 1.31 at −90 mmHg (Figure 3).

## Discussion

Mechanosensitive ion channels have long been suspected to energetically influence each other through cooperative effects. Due to their elastic properties, lipid bilayers propagate membrane deformations induced by distinct channel states, promoting nearby channels to adopt similar conformations. In most cases described in the literature, these inter-channel cooperative effects are mediated by a change of membrane thickness resulting from the hydrophobic mismatch between hydrophobic protein domains and surrounding lipids^24–28^. However, membrane thinning and thickening rapidly decay with lateral distance and most of membrane deformation free energy is brought about by the first annulus of lipids in contact with the protein^29^, thus dramatically limiting the spatial range of cooperative effects. In contrast, our proposed cooperative mechanism occurs through membrane curvature, which is predicted to decay over much longer distances (in typical cell membranes), enabling PIEZO1 channels to potentially influence each other over distances exceeding tens of nanometers^15^.

Our mechanism is consistent with electrophysiology experiments showing collective PIEZO1 gating properties and imaging experiments revealing fluorescently-labeled PIEZO1 channels produce dense fluorescent puncta in cell membranes^17,23,30,31^. PIEZO1 puncta seemingly exhibit a large heterogeneity of size and intensity, suggesting they harbor a variable number of channels. Our mathematical model provides a hypothetical range for the average number of cooperating channels in macroscopic recordings (7 to 15). This number may correspond to the average number of channels per cluster. A more direct experimental approach will be needed to confirm these estimates. Our data further suggests that the free energy change associated with opening of a single channel only decreases by ≈ 3.1 *k*_*B*_T when patch pressure is reduced from 0 to −90 mmHg (assuming 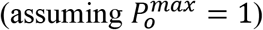). In contrast, increasing the number of channels from 1 to 15 at a constant pressure of −20 mmHg would decrease activation free energy by ≈ 10.5 *kB*T. Thus, the gating free energy of PIEZO1 activation in large channel clusters is predicted to be dominated by cooperative effects rather than membrane tension.

Our mathematical model is likely too simplistic for several reasons. First, our assumption that the activation constant scales by a constant factor each time a channel opens may not capture the complex interplay between membrane energetics and channel diffusion. The cooperativity factor should depend not only on the number of open channels, but also on the average distance between them as channels move laterally in the membrane. Second, our model assumes all channels energetically interact with each other in a membrane patch. However, these channels could be arranged in a variety of cluster configurations that cannot be probed electrophysiologically. While it is reasonable to assume all channels mutually interact within a cluster, it is unclear whether channels from distinct clusters would energetically interact. A deeper understanding of PIEZO1 cooperativity would thus require considering channel diffusion, conformation, membrane footprint energetics and clustering dynamics.

PIEZO1 inactivation is affected by the presence of specific lipids^32,33^, metal ions^30^, disease mutations^34^, as well as external pH^35^, membrane potential^36^, and other unknown cellular factors^37^. In addition to this long list, our study shows that the rate of PIEZO1 inactivation accelerates as more channels open in a patch. Our data also show that the mean open state dwell time lengthens when more channels open. Assuming PIEZO1 inactivation is mediated by stochastic transitions from open to inactivated states, lengthening the duration of opening events would increase the probability of inactivation transitions. The increased surge of ionic flow mediated through cooperative effects may thus be limited by a concomitant acceleration of channel inactivation. Further studies will be needed to probe the link between inter-channel cooperativity and PIEZO1 inactivation.

The modulation of PIEZO1 channels by inter-channel cooperativity has profound physiological implications. This mode of regulation may explain how PIEZO1 channels sense mechanical stimuli across many orders of magnitude, from minute forces induced by capillary lymph flow^38,39^ to stronger forces induced by arterial blood flow and pressure^40–44^. Our data suggest reducing the number of channels per PIEZO1 cluster would reduce cellular mechanosensitivity and *vice versa*. This hypothetical mode of channel regulation would constitute a remarkably effective mechanism to modulate cellular mechanosensitivity without altering the total number of channels at the cell surface, which would require slow and costly membrane trafficking processes. Such a potential regulatory mechanism seems plausible, as a recent study suggests cell migration is accompanied by a dynamic redistribution of endogenous PIEZO1 channels at the cell surface^45^. Future studies will be needed to determine if and how channel density in PIEZO1 clusters changes as a function of physiological and pathological contexts.

Since PIEZO1 and PIEZO2 share the same structure^10–13^, it is tempting to speculate that PIEZO2 channels energetically interact similarly to PIEZO1, a phenomenon that would explain the ability of PIEZO2-dependent mechanoreceptors to sense a large amplitude range of mechanical forces, from gentle touch to large visceral pressures^40,46–48^. However, while both PIEZO1 and PIEZO2 exquisitely respond to positive patch pressurization, PIEZO2 is reportedly weakly sensitive to negative pressurization, suggesting profound differences in tension-sensing mechanism between the two mammalian PIEZO homologs^3,46^. Further experiments are needed to assess whether PIEZO2 channels gate in a cooperative manner.

## Author contributions

T.W, A.D.O and J.J.L conceived the project; T.W. performed electrophysiology experiments; T.W, A.D.O, and J.J.L. analyzed data, A.D.O. performed MATLAB simulations; W.Y and Y.L performed and analyzed molecular dynamics simulations; J.J.L derived equations and wrote the manuscript with input from all authors.

## Acknowledgment

We thank Dr. Medha Pathak for critical reading of the manuscript. This work was supported by NIH grant GM130834 to Y.L and J.J.L.

