## Supplementary Information for "An inter-channel cooperative mechanism mediates PIEZO1’s exquisite mechanosensitivity"

**Supplementary Information for Wijerathne et. al**

This supplementary information contains:

Supplementary Figures 1-9

Supplementary Appendices 1-3

Supplementary References

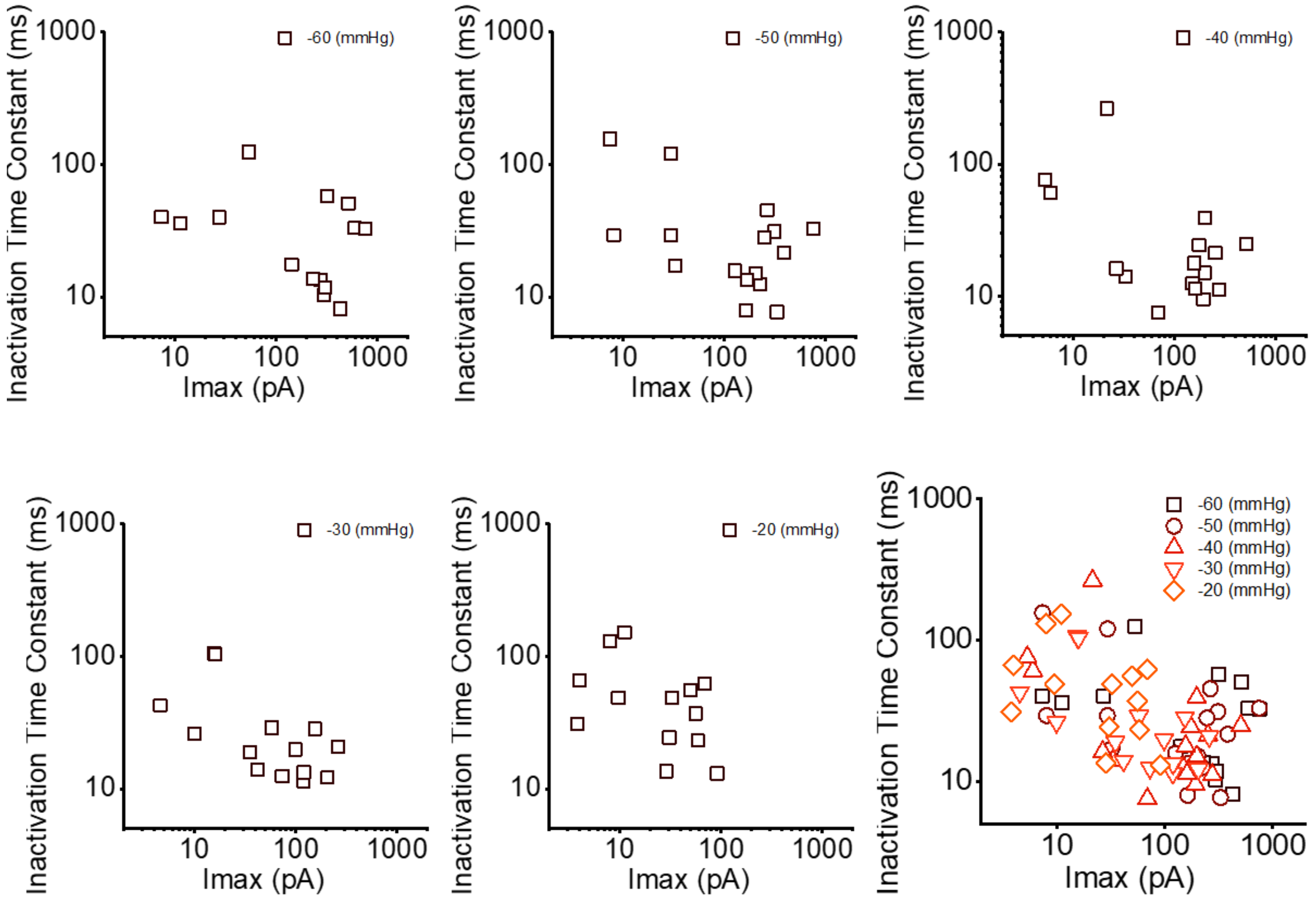


**Supplementary Fig. 1: PIEZO1 inactivation kinetics slows down in patches producing larger saturating currents.** The PIEZO1 inactivation time constant was obtained using a mono-exponential fit of the decaying portion of ionic currents while the pressure pulse was maintained. The plots show the time constant value of individual current trace as a function of the maximal saturating current in the patch ($I_{max}$) and for various pressure pulses from -20 to -60 mmHg. Data are from 16 independent patches. The Pearson's correlation coefficient between Log(inactivation time constant) and Log($I_{max}$) is -0.52.

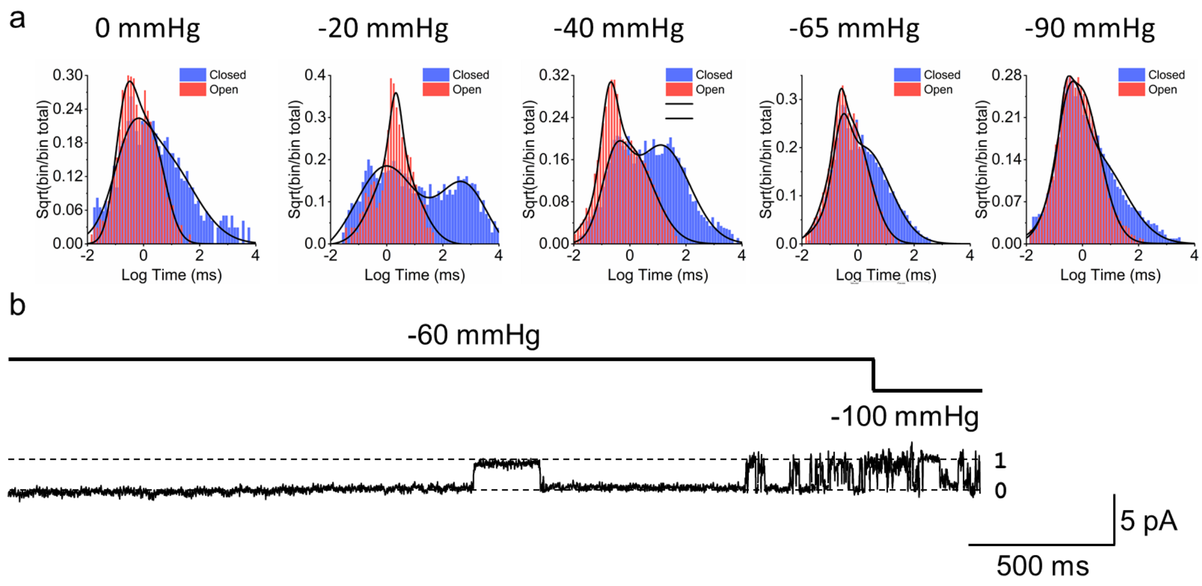


**Supplementary Fig. 2: Analysis of single channel PIEZO1 currents.** (**a**) Dwell time distribution of opening and closing events extracted from current traces using Clampex (Molecular Devices) at the indicated steady-state patch pressure. (**b**) Example of pulse procedure to rule out multichannel traces from single channel analyses. Numbers on the right of the trace indicate the number of open channels. V = -100 mV.

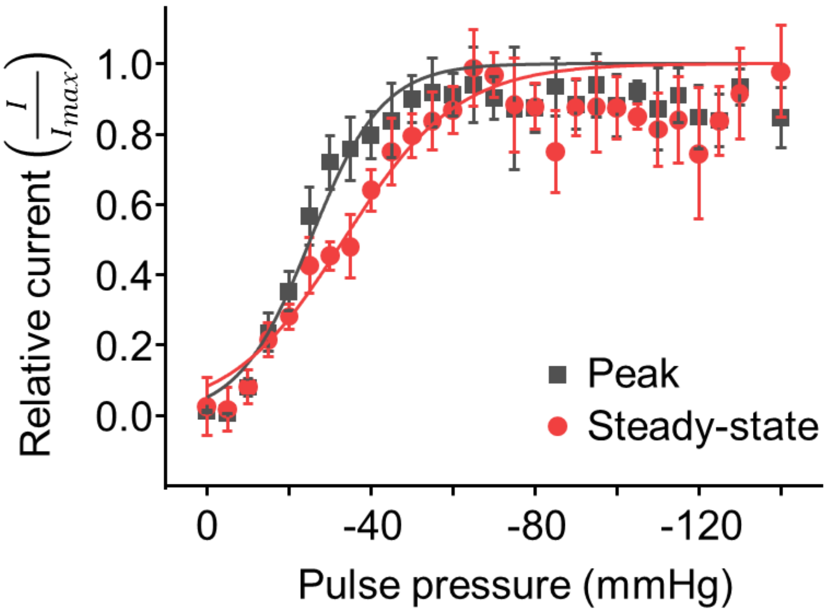


**Supplementary Fig. 3: Pressure-dependence of peak vs. steady-state macroscopic PIEZO1 currents.** Pairwise comparison between relative peak (dark grey) and relative steady-state (red) macroscopic PIEZO1 current as a function of the patch pressure. Curve fitting using equation (1a) gives $p_{1/2}^{app}$ values of -25 ± 2 mmHg (peak) and -33.0 ± 1.5 mmHg (steady-state) and $\varepsilon$ values of 0.117 ± 0.018 $mmHg^{-1}$ (peak) and 0.077 ± 0.009 $mmHg^{-1}$ (steady state). Data are from 11 independent cell-attached patches. Error bars = s.e.m.

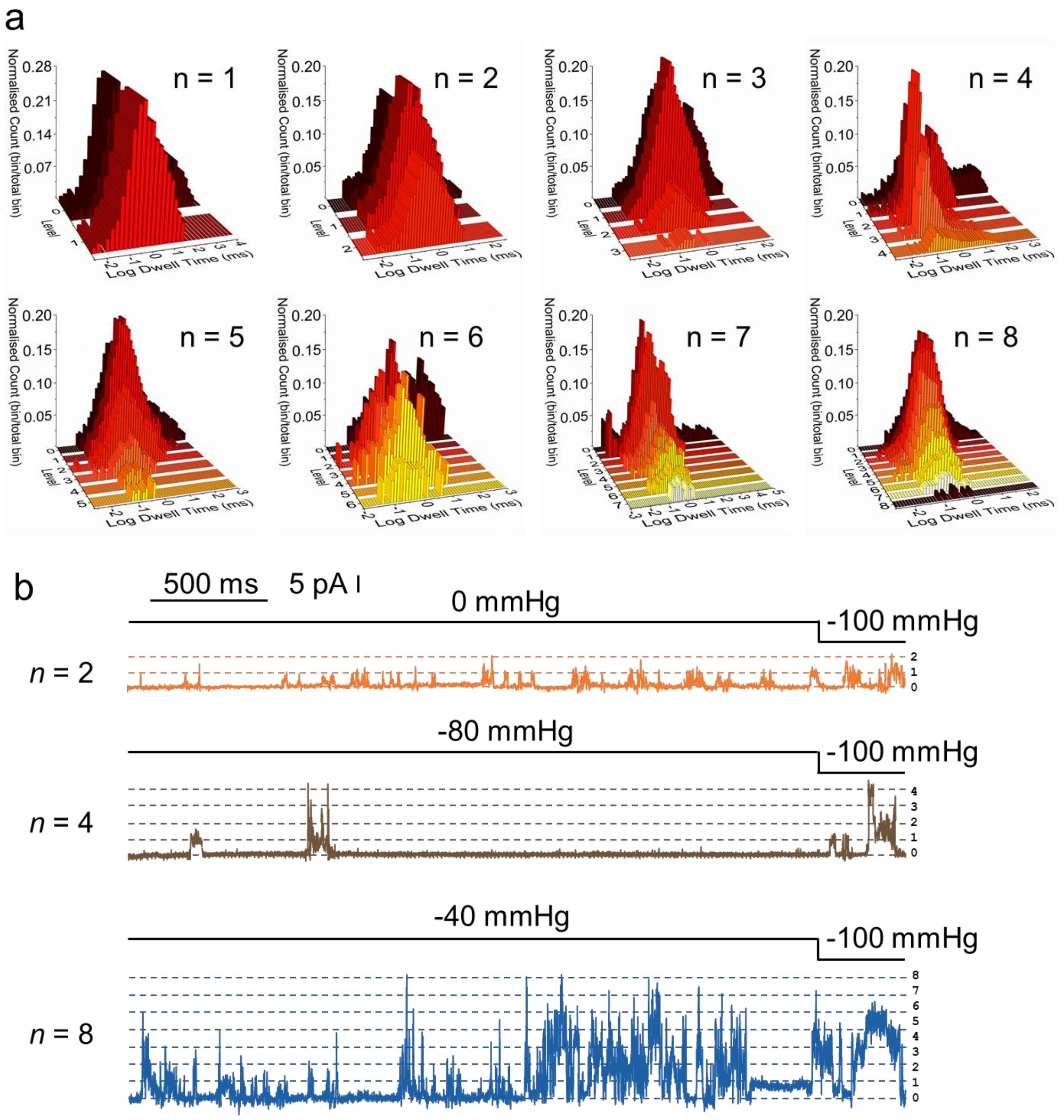


**Supplementary Fig.4: Analysis of multichannel PIEZO1 recordings.** (**a**) Dwell time distribution of multichannel open probabilities from individual multichannel patches pressurized at -40 mmHg (*n* = 1-6, 8) or -90 mmHg (*n* = 7). (**b**) Examples of two-pulse procedure used to confirm the number of channels in multichannel patches. Numbers on the right of the traces represent the number of open channels. V = -100 mV.

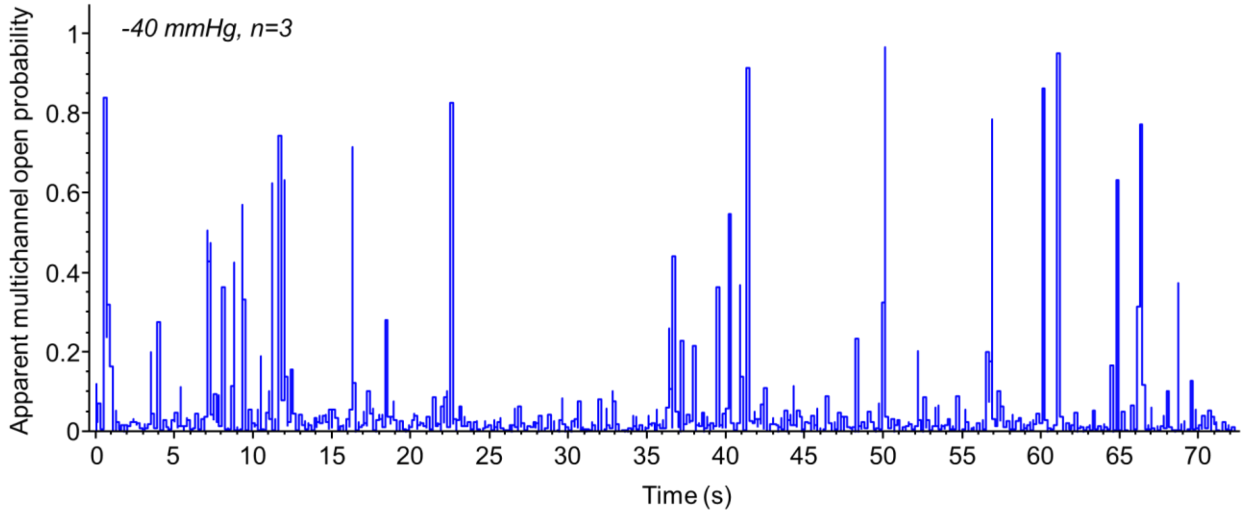


**Supplementary Fig. 5: Time course of steady-state current activity of PIEZO1 channels in a multichannel patch.** The apparent open probability was extracted for a multichannel patch pressurized at -40 mmHg and containing 3 channels (*n*=3) using a 100 ms rolling window (Clampex, Molecular Devices) and plotted as a function the recording time.

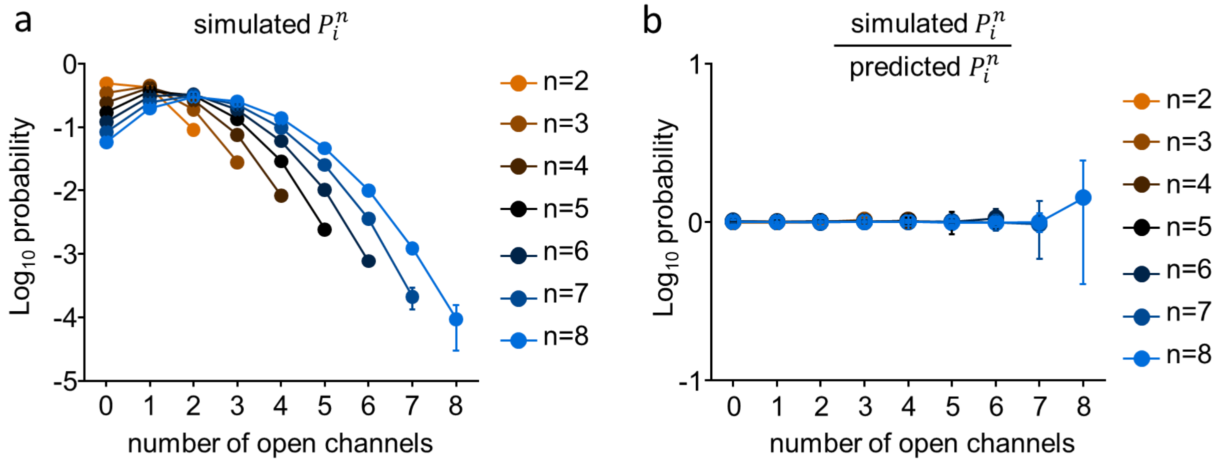


**Supplementary Fig. 6: Simulations of multichannel open probabilities in which channels gate independently.** A MATLAB script was used to simulate stochastic gating transitions in 7 multichannel patches containing 2 to 8 virtual channels with a single channel open probability of 0.3 and a mean open dwell time of 30 ± 10 ms. Each simulated channel underwent 2000 activation events and each simulation was repeated 10 times. (**a**) Plot showing the average $P_i^n$ for all simulations. (**b**) Plots showing the ratio of simulated $P_i^n$ values shown in (a) divided by the $P_i^n$ values calculated using equation (2). Error bars = standard deviations.

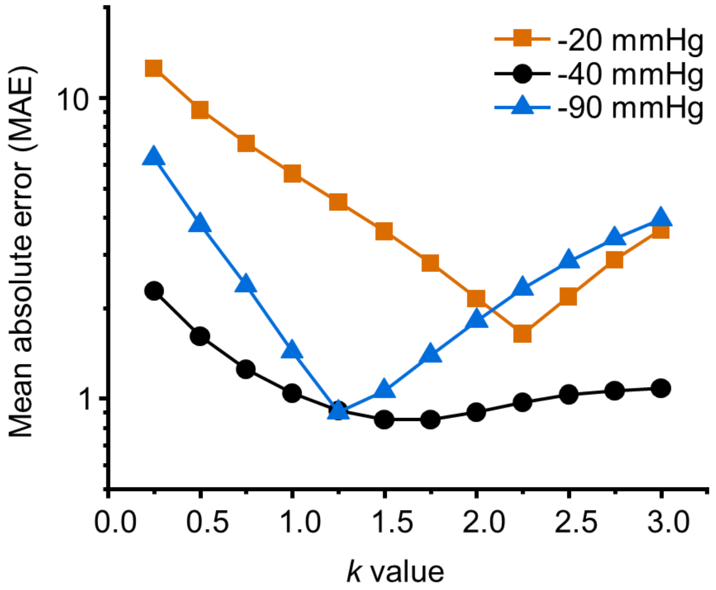


**Supplementary Fig. 7: Fitting the cooperativity parameter *k* through MAE minimization.** MAE values are plotted as a function of the *k* value in the range $0.25 \leq k \leq 3$ for each tested pressure.

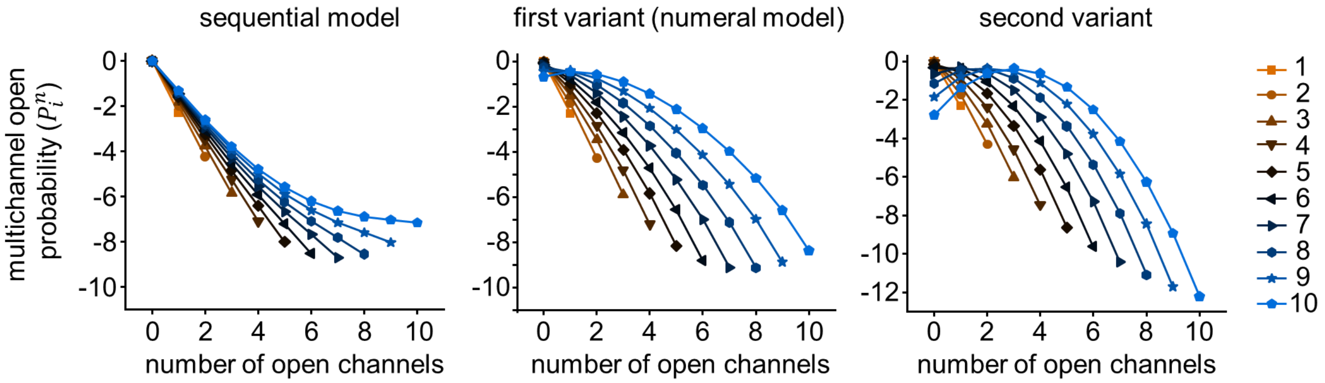


**Supplementary Fig. 8: Predicted $P_i^n$ from the sequential cooperativity model and its two variants.** Predicted multichannel probabilities are calculated from equation (3) (sequential model) or variants of equation (3) in which the exponent of the *k* parameter is set to i(n-1) (first variant) or *i*(2*n*-*i*-1)/2 (second variant). The open probability value was set to 0.005.

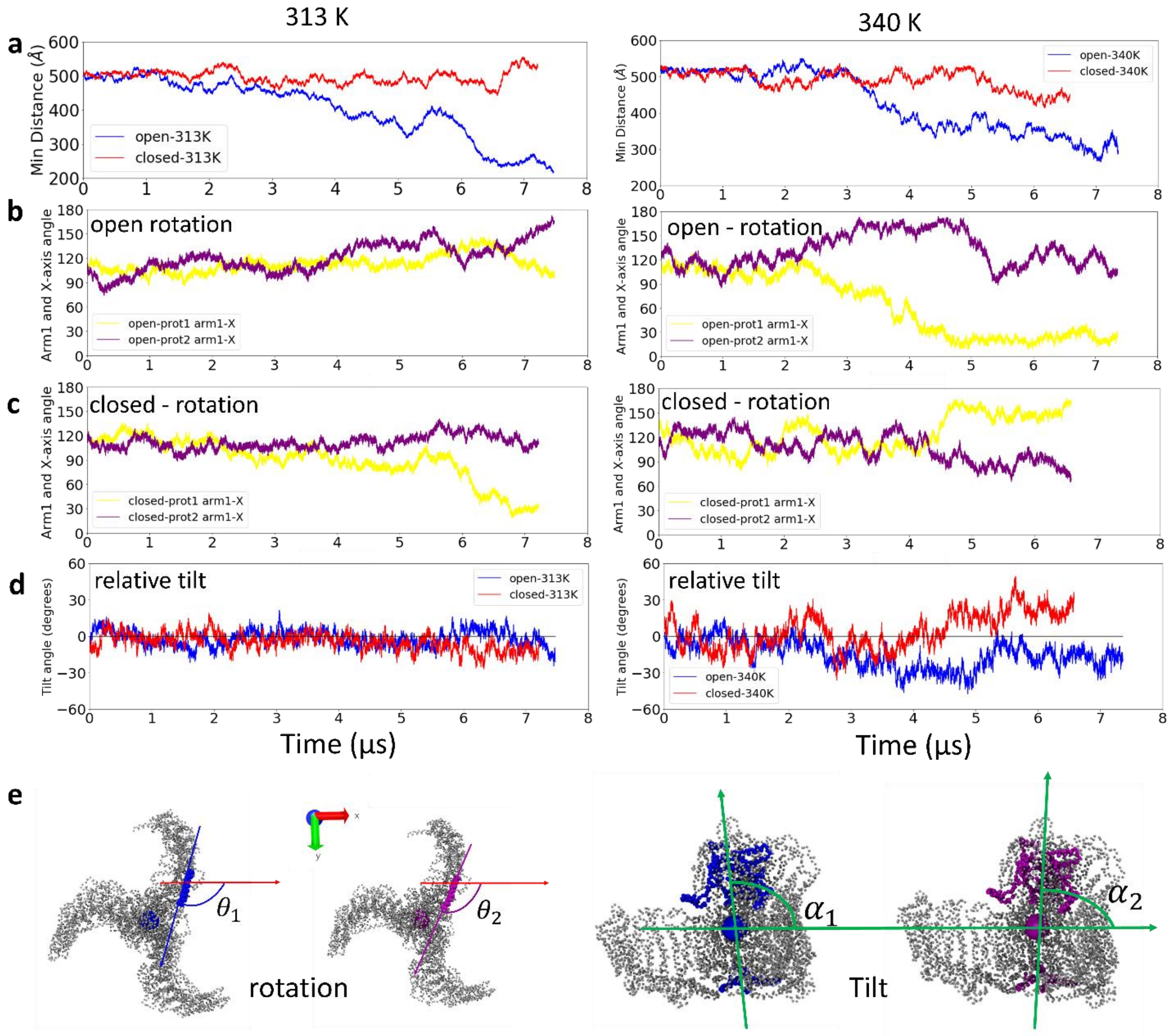


**Supplementary Fig. 9: Minimum pore-pore distance, protein rotation and protein tilting along the simulated trajectories at two temperatures.** (**a**) Time course of minimal pore-pore distance in open-open (blue) and closed-closed (red) simulations. The rotation of each channel determined as the angle between the beam of arm1 in each channel and the x-axis is plotted as function of time for the open-open (**b**) and closed-closed (**c**) systems. (**d**) Relative tilt angle between two proteins (blue for open-open, and red for closed-closed), defined as $\alpha 2 - \alpha 1$. $\alpha$ is the angle between the internal axis of the protein with respect to the horizontal protein-protein vector. The internal axis, which lines the central conduction pathway, is defined as the vector between the intracellular C-terminal domain and the extracellular cap domain. (**e**) The arm rotation ($\theta 1, \theta 2$) and protein tilt angle ($\alpha 2 - \alpha 1$) are illustrated at the bottom.

**Supplementary Appendix 1**

**Free energy activation equations for mechanosensitive PIEZO channels**

The force-from-lipid paradigm posits that the free energy change associated with channel opening, $\Delta G$, relates to membrane tension, $\gamma$, and to the membrane-projected protein expansion area associated with channel opening, $\Delta A_p$:

$$\Delta G = \Delta G_0 - \gamma \Delta A_p \qquad (S1)$$

with $\Delta G_0$ the value of $\Delta G$ in absence of tension[1]. Since PIEZO1 deforms the membrane from its planar shape, a membrane deformation energy term, $\Delta G_m$, must be subtracted to the right side of equation (S1)[2,3]:

$$\Delta G = \Delta G_0 - \Delta G_m - \gamma \Delta A_p$$

The Helfrich's equation[4] indicates that, for a homogenous fluid-like membrane, $G_m$ can be approximated by the sum of two energy components:

$$G_m^{bending} = \frac{1}{2} K_c \int (c_1 + c_2)^2 \, dA$$

$$G_m^{stretching} = \gamma A_m$$

with $K_c$, the membrane bending modulus, $c_1$ and $c_2$ the principal curvatures of mid-bilayer surface $A$, and $A_m$, the in-plane membrane surface expansion due to its deformation away from a planar surface. This leads to a more comprehensive expression of the free energy activation for a single PIEZO channel:

$$\Delta G = \Delta G_0 - \Delta G_m^{bending} - \gamma \left( \Delta A_m + \Delta A_p \right)$$

This expression simplifies to:

$$\Delta G = (\gamma_{1/2} - \gamma)(\Delta A_m + \Delta A_p) \qquad (S2)$$

with $\gamma_{1/2}$, the tension at which $\Delta G = 0$. The activation free energy for single channels can be obtained from the Boltzmann definition of free energy:

$$\Delta G = -k_B T \, ln\left(\frac{\alpha_1^1}{\beta_1^1}\right) = -k_B T \, ln\left(\frac{P_o/P_o^{max}}{1 - P_o/P_o^{max}}\right) \qquad (S3)$$

Although increasing tension necessarily favors the flatter channel conformation, it is unknown whether the maximal channel open probability induced by tension equates 1, hence the $P_o/P_o^{max}$ term.

By combining equations (S2) and (S3):

$$-k_B T \, ln\left(\frac{P_o/P_o^{max}}{1 - P_o/P_o^{max}}\right) = (\gamma_{1/2} - \gamma)(\Delta A_m + \Delta A_p)$$

$$\Leftrightarrow \quad \frac{P_o/P_o^{max}}{1 - P_o/P_o^{max}} = e^{-\left(\frac{(\gamma_{1/2}-\gamma)(\Delta A_m+\Delta A_p)}{k_B T}\right)}$$

$$\Leftrightarrow \quad P_o/P_o^{max} = (1 - P_o/P_o^{max})e^{-\left(\frac{(\gamma_{1/2}-\gamma)(\Delta A_m+\Delta A_p)}{k_B T}\right)}$$

$$\Leftrightarrow \quad P_o/P_o^{max}\left[1 + e^{-\left(\frac{(\gamma_{1/2}-\gamma)(\Delta A_m+\Delta A_p)}{k_B T}\right)}\right] = e^{-\left(\frac{(\gamma_{1/2}-\gamma)(\Delta A_m+\Delta A_p)}{k_B T}\right)}$$

After simplification:

$$P_o/P_o^{max} = \frac{1}{1 + e^{\left(\frac{(\gamma_{1/2}-\gamma)(\Delta A_m+\Delta A_p)}{k_B T}\right)}}$$

According to the Young-Laplace law, we have:

$$p = -\frac{2\gamma}{R}$$

With $R$ the radius of membrane curvature produced by pipette aspiration and $p$ a negative pressure pulse applied to the pipette. For identical pipette geometry (resistance and tip diameter), the membrane radius $R$ and the corresponding tension $\gamma$ are proportional to the pressure gradient $p$. Therefore, we can write:

$$P_o = \frac{P_o^{max}}{1 + e^{(p-p_{1/2})\varepsilon}} \qquad (1b)\ in\ the\ main\ text$$

With $p_{1/2}$, the pressure producing $P_o = 0.5$, and $\varepsilon$, a slope factor equal to $\frac{(\Delta A_m+\Delta A_p)}{k_B T}$.

This equation is often used to fit the apparent open probability from macroscopic recordings:

$$\frac{I}{I_{max}} = \frac{1}{1 + e^{(p-p_{1/2}^{app})\varepsilon}} \qquad (1a)\ in\ the\ main\ text$$

#### Supplementary Appendix 2

##### 1. Equations to predict open probabilities for non-independent gating (sequential model)

Let us consider a cluster containing a single channel in equilibrium between a conducting (open, O) and non-conducting (closed, C) state according to rate constants α and β:

$$C \underset{\beta}{\overset{\alpha}{\rightleftarrows}} O$$

As for any chemical transformation, the equilibrium constant, α/β, is equal to the concentration ratio of open/closed species:

$$\frac{[O]}{[C]} = \frac{\alpha}{\beta}$$

Let us define $P_i^n$ as the probability of a cluster harboring n channels to have i channel(s) open (i ≤ n). Using this terminology, the single channel open probability is $P_o$, and the closed probability is $1 - P_o$, with $P_o = P_1^1$. For a cluster containing a single channel, we have:

$$P_o = \frac{[O]}{[O] + [C]} = \frac{\alpha}{\alpha + \beta}$$

$$1 - P_o = \frac{[C]}{[O] + [C]} = \frac{\beta}{\alpha + \beta}$$

Thus, we can write:

$$\frac{\alpha}{\beta} = \frac{P_o}{1 - P_o}$$

Now, let us consider the linear scheme below representing gating transitions in a cluster of 5 identical channels which gate independently. Since there are multiple ways to populate the intermediate macroscopic conductance levels with 1, 2, 3, or 4 channels open, the opening and closure rate constants are multiplied by a coefficient representing the number of possible microscopic transitions for each step:

$$CCCCC \underset{\beta}{\overset{5\alpha}{\rightleftarrows}} CCCCO \underset{2\beta}{\overset{4\alpha}{\rightleftarrows}} CCCOO \underset{3\beta}{\overset{3\,\alpha}{\rightleftarrows}} CCOOO \underset{4\beta}{\overset{2\alpha}{\rightleftarrows}} COOOO \underset{5\beta}{\overset{\alpha}{\rightleftarrows}} OOOOO$$

Let us use "$L_i$" to denote the conductance level with *i* channel(s) open:

$$L_0 \underset{\beta_1^5}{\overset{5\alpha_1^5}{\rightleftarrows}} L_1 \underset{2\beta_2^5}{\overset{4\alpha_2^5}{\rightleftarrows}} L_2 \underset{3\beta_3^5}{\overset{3\,\alpha_3^5}{\rightleftarrows}} L_3 \underset{4\beta_4^5}{\overset{2\alpha_4^5}{\rightleftarrows}} L_4 \underset{5\beta_5^5}{\overset{\alpha_5^5}{\rightleftarrows}} L_5$$

Under our assumption for cooperative gating, the microscopic equilibrium constant, $\frac{\alpha}{\beta}$ , scales by $k$ for each iteration of $i$. In this "sequential model", we have:

$$\frac{\alpha_{i+1}^n}{\beta_{i+1}^n} = k\frac{\alpha_i^n}{\beta_i^n} \quad \Leftrightarrow \quad \frac{\alpha_i^n}{\beta_i^n} = k^{(i-1)}\frac{\alpha_1^n}{\beta_1^n}$$

Let us now express the concentrations of all possible macroscopic configurations as a function of $[L_0]$.

<u>1 channel open:</u>

$$[L_1] = 5\frac{\alpha_1^5}{\beta_1^5}[L_0]$$

<u>2 channels open:</u>

$$[L_2] = \frac{4}{2}\frac{\alpha_2^5}{\beta_2^5}[L_1] = 2\frac{\alpha_2^5}{\beta_2^5}\ 5\ \frac{\alpha_1^5}{\beta_1^5}[L_0] = 10k\ \frac{\alpha_1^5}{\beta_1^5}\ \frac{\alpha_1^5}{\beta_1^5}\ [L_0] = 10\ k\ \left(\frac{\alpha_1^5}{\beta_1^5}\right)^2\ [L_0]$$

<u>3 channels open:</u>

$$[L_3] = \frac{3}{3}\frac{\alpha_3^5}{\beta_3^5}[L_2] = k^2\frac{\alpha_1^5}{\beta_1^5}10\ k\left(\frac{\alpha_1^5}{\beta_1^5}\right)^2\ [L_0] = 10\ k^3\left(\frac{\alpha_1^5}{\beta_1^5}\right)^3\ [L_0]$$

<u>4 channels open:</u>

$$[L_4] = \frac{2}{4}\frac{\alpha_4^5}{\beta_4^5}[L_3] = \frac{2}{4}\frac{\alpha_4^5}{\beta_4^5}10\ k^3\left(\frac{\alpha_1^5}{\beta_1^5}\right)^3\ [L_0] = 5k^3\frac{\alpha_1^5}{\beta_1^5}\ k^3\left(\frac{\alpha_1^5}{\beta_1^5}\right)^3\ [L_0] = 5k^6\left(\frac{\alpha_1^5}{\beta_1^5}\right)^4\ [L_0]$$

<u>5 channels open:</u>

$$[L_5] = \frac{1}{5}\frac{\alpha_5^5}{\beta_5^5}[L_4] = k^4\frac{\alpha_1^5}{\beta_1^5}\ k^6\left(\frac{\alpha_1^5}{\beta_1^5}\right)^4\ [L_0] =\ k^{10}\left(\frac{\alpha_1^5}{\beta_1^5}\right)^5\ [L_0]$$

Note that the exponent of the ratio $\frac{\alpha_i^n}{\beta_i^n}$ in these expressions is equal to $i$. In addition, the positive integer in each expression represents a binomial coefficient. Also, note that the exponent of the cooperativity coefficient, $a_i$, is the arithmetic series of the finite arithmetic progression:

$$a_i = i - 1$$

with i ≥ 1. The Gauss's formula equates the arithmetic series, $S$, to:

$$S = \sum_{i=1}^{i} a_i = \frac{(i-1)i}{2}$$

Note that this number corresponds to the number of order-independent permutations without repetitions of pairs of open channels when *i* open channels are present.

According to the assumptions for this model, we have:

$$\frac{\alpha_1^n}{\beta_1^n} = \frac{\alpha_1^1}{\beta_1^1} = \frac{P_o}{1-P_o}$$

This leads to the general expression:

$$[L_i^n] = \binom{n}{i} k^{\frac{(i-1)i}{2}} \left(\frac{P_o}{1-P_o}\right)^i [L_0]$$

Now, we can express the probability of each $L_i^n$ level for any value of *i* and *n*. As an example, let us first calculate the probability of having one open channel in the cluster of 5 channels, $P_1^5$:

$$P_1^5 = \frac{[L_1]}{[L_0]+[L_1]+[L_2]+[L_3]+[L_4]+[L_5]}$$

$$P_1^5 = \frac{\binom{5}{1}\left(\frac{P_o}{1-P_o}\right)^1 [L_0]}{[L_0] + \binom{5}{1}\left(\frac{P_o}{1-P_o}\right)^1 [L_0] + \binom{5}{2} k^1 \left(\frac{P_o}{1-P_o}\right)^2 [L_0] + \binom{5}{3} k^3 \left(\frac{P_o}{1-P_o}\right)^3 [L_0] + \binom{5}{4} k^6 \left(\frac{P_o}{1-P_o}\right)^4 [L_0] + \binom{5}{5} k^{10} \left(\frac{P_o}{1-P_o}\right)^5 [L_0]}$$

Dividing each term by $[L_0]$:

$$P_1^5 = \frac{5k^0\left(\frac{P_o}{1-P_o}\right)^1}{1 + 5\left(\frac{P_o}{1-P_o}\right)^1 + 10\,k^1\left(\frac{P_o}{1-P_o}\right)^2 + 10\,k^3\left(\frac{P_o}{1-P_o}\right)^3 + 5k^6\left(\frac{P_o}{1-P_o}\right)^4 + k^{10}\left(\frac{P_o}{1-P_o}\right)^5}$$

By generalization, we have:

$$P_i^n = \frac{\binom{n}{i} k^{\binom{i}{2}} \left(\frac{P_o}{1-P_o}\right)^i}{\sum_{i=0}^{n} \binom{n}{i} k^{\binom{i}{2}} \left(\frac{P_o}{1-P_o}\right)^i} \qquad \textit{equation (3) in the main text}$$

Note that when $k = 1$, equation (3) is equivalent to:

$$P_i^n = \frac{\binom{n}{i} \left(\frac{P_o}{1-P_o}\right)^i}{\sum_{i=0}^{n} \binom{n}{i} \left(\frac{P_o}{1-P_o}\right)^i}$$

Which is equivalent to the combinatorial expression for independent gating:

$$P_i^n = \binom{n}{i} (P_o)^i (1-P_o)^{(n-i)} \qquad \textit{equation (2) in the main text}$$

<u>Proof:</u>

$$\binom{n}{i} (P_o)^i (1-P_o)^{(n-i)} = \binom{n}{i} \left(P_o \frac{1-P_o}{1-P_o}\right)^i (1-P_o)^{(n-i)} = \binom{n}{i} \left(\frac{P_o}{1-P_o}\right)^i (1-P_o)^n$$

Thus,

$$\binom{n}{i} (P_o)^i (1-P_o)^{(n-i)} = \frac{\binom{n}{i} \left(\frac{P_o}{1-P_o}\right)^i}{\sum_{i=0}^{n} \binom{n}{i} \left(\frac{P_o}{1-P_o}\right)^i} \Leftrightarrow \binom{n}{i} \left(\frac{P_o}{1-P_o}\right)^i (1-P_o)^n = \frac{\binom{n}{i} \left(\frac{P_o}{1-P_o}\right)^i}{\sum_{i=0}^{n} \binom{n}{i} \left(\frac{P_o}{1-P_o}\right)^i}$$

Dividing each side by $\binom{n}{i} \left(\frac{P_o}{1-P_o}\right)^i$:

$$\Leftrightarrow (1-P_o)^n = \frac{1}{\sum_{i=0}^{n} \binom{n}{i} \left(\frac{P_o}{1-P_o}\right)^i}$$

$$\Leftrightarrow (1-P_o)^n = \frac{1}{\sum_{i=0}^{n}\binom{n}{i}(P_o)^i(1-P_o)^{-i}}$$

$$\Leftrightarrow (1-P_o)^n \sum_{i=0}^{n}\binom{n}{i}(P_o)^i(1-P_o)^{-i} = 1$$

$$\Leftrightarrow \sum_{i=0}^{n}\binom{n}{i}(P_o)^i(1-P_o)^{(n-i)} = 1$$

The left side term of this equation is the sum of the probabilities of all possible macroscopic configurations of the system. This sum is necessarily equal to unity, i.e. the right-side term.

**2. Equations to predict open probabilities for non-independent gating (numeral model)**

Another assumption for cooperative gating posits that the microscopic equilibrium constant, $\frac{\alpha}{\beta}$, scales by *k* for each iteration of *n*. In this "numeral model", we have:

$$\frac{\alpha_i^{n+1}}{\beta_i^{n+1}} = k\frac{\alpha_i^n}{\beta_i^n} \quad \Leftrightarrow \quad \frac{\alpha_i^n}{\beta_i^n} = k^{(n-1)}\frac{\alpha_1^1}{\beta_1^1}$$

Let us quantify the concentrations of clusters with i channel open and with various n

<u>1 channel:</u>

$$[L_1] = \frac{\alpha_i^1}{\beta_i^1}[L_0]$$

$$P_1^1 = \frac{\frac{\alpha_i^1}{\beta_i^1}}{1+\frac{\alpha_i^1}{\beta_i^1}}$$

<u>2 channels:</u>

$$[L_1] = 2\frac{\alpha_1^2}{\beta_1^2}[L_0] = \ 2k\frac{\alpha_1^1}{\beta_1^1}[L_0]$$

$$[L_2] = \frac{1}{2}\frac{\alpha_2^2}{\beta_2^2}[L_1] = \frac{1}{2}\frac{\alpha_2^2}{\beta_2^2}\ 2k\frac{\alpha_1^1}{\beta_1^1}[L_0] = k^2\left(\frac{\alpha_1^1}{\beta_1^1}\right)^2\ [L_0]$$

$$P_1^2 = \frac{2k\frac{\alpha_i^1}{\beta_i^1}}{1 + 2k\frac{\alpha_i^1}{\beta_i^1} + k^2\left(\frac{\alpha_i^1}{\beta_i^1}\right)^2}$$

$$P_2^2 = \frac{k^2\left(\frac{\alpha_i^1}{\beta_i^1}\right)^2}{1 + 2k\frac{\alpha_i^1}{\beta_i^1} + k^2\left(\frac{\alpha_i^1}{\beta_i^1}\right)^2}$$

<u>3 channels:</u>

$$[L_1] = 3\frac{\alpha_1^3}{\beta_1^3}[L_0] = 3k^2\frac{\alpha_1^1}{\beta_1^1}[L_0]$$

$$[L_2] = \frac{\alpha_2^3}{\beta_2^3}[L_1] = \frac{\alpha_2^3}{\beta_2^3}3k^2\frac{\alpha_1^1}{\beta_1^1}[L_0] = k^2\frac{\alpha_1^1}{\beta_1^1}3k^2\frac{\alpha_1^1}{\beta_1^1}[L_0] = 3k^4\left(\frac{\alpha_1^1}{\beta_1^1}\right)^2[L_0]$$

$$[L_3] = \frac{1}{3}\frac{\alpha_3^3}{\beta_3^3}[L_2] = \frac{1}{3}\frac{\alpha_3^3}{\beta_3^3}3k^4\left(\frac{\alpha_1^1}{\beta_1^1}\right)^2[L_0] = k^2\frac{\alpha_1^1}{\beta_1^1}k^4\left(\frac{\alpha_1^1}{\beta_1^1}\right)^2[L_0] = k^6\left(\frac{\alpha_1^1}{\beta_1^1}\right)^3[L_0]$$

$$P_1^3 = \frac{3k^2\frac{\alpha_1^1}{\beta_1^1}}{1 + 3k^2\frac{\alpha_1^1}{\beta_1^1} + 3k^4\left(\frac{\alpha_1^1}{\beta_1^1}\right)^2 + k^6\left(\frac{\alpha_1^1}{\beta_1^1}\right)^3}$$

$$P_2^3 = \frac{3k^4\left(\frac{\alpha_1^1}{\beta_1^1}\right)^2}{1 + 3k^2\frac{\alpha_1^1}{\beta_1^1} + 3k^4\left(\frac{\alpha_1^1}{\beta_1^1}\right)^2 + k^6\left(\frac{\alpha_1^1}{\beta_1^1}\right)^3}$$

$$P_3^3 = \frac{k^6\left(\frac{\alpha_1^1}{\beta_1^1}\right)^3}{1 + 3k^2\frac{\alpha_1^1}{\beta_1^1} + 3k^4\left(\frac{\alpha_1^1}{\beta_1^1}\right)^2 + k^6\left(\frac{\alpha_1^1}{\beta_1^1}\right)^3}$$

A generalization to *n* channels leads to:

$$P_i^n = \frac{\binom{n}{i}k^{(n-1)i}\left(\frac{P_1^1}{1-P_1^1}\right)^i}{\sum_{i=0}^{n}\binom{n}{i}k^{(n-1)i}\left(\frac{P_1^1}{1-P_1^1}\right)^i}$$

### Supplementary Appendix 3

**Determination of macroscopic $\frac{I}{I_{max}}$ from single channel parameters under the assumption of sequential cooperativity**

The total current *I* at time *t* is a function of the number of open channels at this instant. In a patch with *n* channels, the current is maximal when *n* channels are open, i.e. when $P_{i=n}^{n} = 1$. If only one channel is open, the current is equal to $\frac{I_{max}}{n}$. If the probabilities for each conductance level are known, the $\frac{I}{I_{max}}$ values can be calculated by first multiplying each conductance level probability by the corresponding number of open channels (*i*), taking the sum of these probabilities, and dividing it by *n*:

$$\frac{I}{I_{max}} = \frac{1}{n}\left\{\frac{[L_1] + 2[L_2] + 3[L_3] + \cdots + n[L_n]}{[L_0] + [L_1] + [L_2] + [L_3] + \cdots + [L_n]}\right\}$$

$$\Leftrightarrow \frac{I}{I_{max}} = \frac{1}{n}\sum_{i=0}^{n} i\ P_i^n$$

Replacing $P_i^n$ values by their expression under the assumption of sequential cooperativity, we have:

$$\frac{I}{I_{max}} = \frac{1}{n}\frac{\sum_{i=0}^{n}\binom{n}{i}\, i\, k^{\binom{i}{2}}\left(\frac{P_o}{1-P_o}\right)^i}{\sum_{i=0}^{n}\binom{n}{i}\ k^{\binom{i}{2}}\left(\frac{P_o}{1-P_o}\right)^i}$$

$\left(\frac{P_o}{1-P_o}\right)$ can be expressed as function of the single channel gating free energy $\Delta G$:

$$\frac{I}{I_{max}} = \frac{1}{n}\frac{\sum_{i=0}^{n}\binom{n}{i}\, i\, k^{\binom{i}{2}}\left(e^{(p_{1/2}-p)\varepsilon}\right)^i}{\sum_{i=0}^{n}\binom{n}{i}\ k^{\binom{i}{2}}\left(e^{(p_{1/2}-p)\varepsilon}\right)^i}$$

After simplification:

$$\frac{I}{I_{max}} = \frac{\sum_{i=0}^{n}\binom{n}{i}\, i\, k^{\binom{i}{2}}\left(e^{(p_{1/2}-p)\varepsilon}\right)^i}{n\sum_{i=0}^{n}\binom{n}{i}\ k^{\binom{i}{2}}\left(e^{(p_{1/2}-p)\varepsilon}\right)^i} \qquad \textit{equation (4) in the main text}$$
